# Criticality and network structure drive emergent oscillations in a stochastic whole-brain model

**DOI:** 10.1101/2022.01.17.476567

**Authors:** Giacomo Barzon, Giorgio Nicoletti, Benedetta Mariani, Marco Formentin, Samir Suweis

## Abstract

Understanding the relation between the structure of brain networks and its functions is a fundamental open question. Simple models of neural activity based on real anatomical networks have proven to be effective in describing features of whole-brain spontaneous activity when tuned at their critical point. In this work, we show that indeed structural networks are a crucial ingredient in the emergence of collective oscillations in a whole-brain stochastic model at criticality. We study analytically a stochastic Greenberg-Hastings cellular automaton in the mean-field limit, showing that it undergoes an abrupt phase transition with a bistable region. In particular, no global oscillations emerge in this limit. Then, we show that by introducing a network structure in the homeostatic normalization regime, the bistability may be disrupted, and the transition may become smooth. Concomitantly, through an interplay between the network topology and weights, a large peak in the power spectrum appears around the transition point, signaling the emergence of collective oscillations. Hence, both the structure of brain networks and criticality are fundamental in driving the collective responses of whole-brain stochastic models.

## 1. Introduction

The human brain is an impressively complex system, spanning several spatial scales of organizations, from microcircuits to whole-brain networks. The comprehensive map of neural connections is usually referred to as “connectome” [1]. However, it is typically unfeasible to reconstruct connectomes at the neuronal level, and often one relies on anatomical connectivity at coarser spatial scales. In humans, such brain structural networks are typically assessed with diffusion tensor/spectrum imaging techniques, which quantify the white matter pathways between mesoscopic brain regions [2, 3].

These complex interconnections act as a backbone on top of which the neurophysiological dynamics occurs. One way to measure such neural activity is through functional magnetic resonance imaging (fMRI). Correlations in the fMRI signals of spontaneous activity during rest have been repeatedly observed [4], yielding detailed maps of complex emergent patterns of coherent brain activities, called resting state (functional) networks (RSN) [5]. Such patterns, consistent among healthy individuals [6], are specifically associated with neuronal systems responsible for sensory, cognitive, and behavioral functions [7, 8].

A hypothesis that is increasingly being considered in light of growing experimental [9, 10] and theoretical [11, 12] results is that collective emergent patterns are signatures of brain self-organization to a *critical* point [13, 14], i.e., the brain dynamics may be poised at the edge of a phase transition. Over the years, evidence to support this hypothesis emerged in the presence of scale-free neural avalanches [15] and cluster size distributions [16, 17], long-range temporal and spatial [18, 19] correlations during spontaneous brain activity - exemplary properties of a system near its critical point. Furthermore, it was recently shown that the collective dynamics of neurons may be associated with a non-trivial fixed point of phenomenological renormalization groups [20, 21]. Some works have also suggested that such phenomenology is compatible with systems between an asynchronous and a synchronous phase, with emerging oscillations [22, 23, 24]. In all these studies the role of the network structure in driving such emerging patterns - e.g., global oscillations or optimal information processing - is often missing.

In fact, the emerging collective dynamics in the brain is shaped both by the underlying connectome and by the neural population activities [25, 26, 27]. Despite a direct relation between structural and functional networks, to what extent structure does determine the neural dynamics and its critical signatures has still to be clarified [28, 29]. Computational models may be the key to bridging this gap [30]. To this end, biophysically inspired models of neural dynamics are typically built on top of empirically derived structural networks, with the aim of reconciling functional behavior.

Notably, a stochastic version of the Greenberg & Hastings (GH) cellular automaton [31] - one of the simplest models to describe the neural dynamics - running over a human connectome of *N* = 998 cortical regions [32] was shown to match some features of whole-brain activity when tuned to the critical point [16, 18]. Indeed, the model undergoes a critical percolation-like transition in the sizes of active clusters, as a function of the level of induced excitatory activation by neighboring neurons. Yet, it is known that geometrical percolation transitions may arise in stochastic dynamical systems, and they usually do not coincide with actual dynamical transitions [33]. In fact, a dynamical transition that separates a regime of low activity from an over-active phase is present beyond the static percolation transition, and less is known about it. Recent numerical studies have suggested that it may be continuous for certain levels of connectivity, otherwise being discontinuous or even absent [34]. Nevertheless, the mechanisms underlying such transition and a corresponding analytical description of it is still lacking. Here, we will focus on the presence and the properties of such dynamical transition, so to better elucidate the relation between the network structure, brain criticality, and emergent collective oscillations.

To this aim, we develop a stochastic continuous-time formulation of the GH model via a master equation approach. We show analytically how two stable equilibria emerge in the mean-field limit, together with a bistable region of the parameter space where these two equilibria coexist. Hence, the mean field limit predicts a discontinuous transition - i.e., a transition in which the order parameters displays a finite jump. Then, we derive the power spectrum of the oscillations and we show that in general, in such mean-field limit, no characteristic peak is present, that is, we do not observe neural activity with collective oscillations. However, when we go beyond the mean-field by adding a network connecting different brain regions, the picture is quite different. We find that the transition becomes continuous - i.e., the order parameter changes smoothly - and collective sustained oscillations emerge.

Overall, our results shed light on the role of the underlying network structure in the emergent collective patterns observed in the brain, as well as explain the mechanisms behind the phase diagram of the Greenberg & Hastings model reported in previous works [16, 18, 34, 35, 36, 37].

## 2. Methods

### 2.1. Whole-brain stochastic continuous-time model

Here, we develop a continuous-time formulation of the whole brain stochastic model introduced by Haimovici *et al*. [18] to describe the dynamics of the human brain at a mesoscopic scale. Such a model is a variation of the Greenberg & Hastings cellular automaton [31], originally designed to study excitable media. Briefly, each node in the system belongs to one of three states: quiescent *Q*, excited *E*, or refractory *R*. The original dynamics of the GH automaton is modified in such a way that the states undergo the following stochastic transitions:

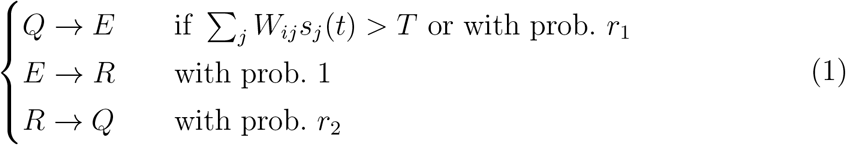

where *s_j_*(*t*) ∈ {0, 1} is the state of node *j* at a certain time step *t* - set to 1 if the node is in the *E* state, and 0 otherwise -, *W_ij_* is the weighted connectivity matrix of the underlying network, *r*_1_ is the probability of self-activation and *r*_2_ is the probability of recovery from the refractory state. In particular, *T* is a threshold governing the induced activation due to interaction with neighboring nodes, which acts as a control parameter of the model.

Hence, in this model, a neuron may either be activated if the weighted combined activity of neighboring neurons exceeds a threshold *T*, or it may self-activate with a probability *r*_1_ that encodes, e.g., external stimuli or unobserved pathways. After activation, neurons switch to a refractory state with unitary probability and cannot activate again. Finally, the escape from the refractory state happens with probability *r*_2_. In this formulation, the state of the system evolves in discrete time steps and is updated synchronously. In particular, for small values of *T* the activity spreads easily between neighboring nodes, even along weak connections. This leads to a regime of high and sustained activation, characterized by fast and temporally uncorrelated fluctuations. We refer to this phase as “super-critical”. For high values of *T*, the activity is instead sustained only by few strong connections, resulting in a suppressed or “sub-critical” phase with regular, short-propagating activity in which nodes fail to give rise to relevant patterns of activity. Importantly, we include homeostatic plasticity in the model, implemented as a normalization of the incoming node’s excitatory input. It has been shown that its addition improves the correspondence between simulated neural patterns and experimental brain functional data [16].

We now study analytically its continuous time, mean-field behavior in the large *N* limit, together with its power spectrum in the stochastic linearized regime. Given a network of *N* units, we denote by *σ_i_*(*t*) ∈ {*E, R, Q*}, *i* = 1,…, *N*, the state of the site *i* at time *t*. The dynamics in (1) can be translated into the following continuous-time evolution: for *h* > 0 and each node i, the probability of having *σ_i_*(*t* + *h*) = *E* given that *σ_i_*(*t*) = *Q* is *r*_act_(*i*)*h* + *o*(*h*) where *r*_act_(*i*) is the rate of activation, defined as

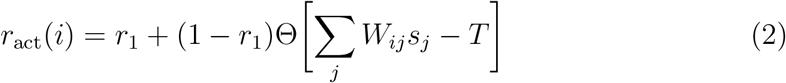

with Θ[·] the Heaviside step function. Notice that 0 ≤ *r*_1_ ≤ 1 by construction. In a similar manner the probability of jumping from state *E* at time *t* to state *R* at time *t* + *h* will be *h* + *o*(*h*) and from *R* to *Q* will be *r*_2_*h* + *o*(*h*) ‡.

The mean-field approximation of the model corresponds to the assumption that the underlying graph is fully-connected with constant weights, i.e., *W_ij_* = *c*, ∀*i, j*. In fact, considering the homeostatic normalization [16], the weights of the structural matrix are simply 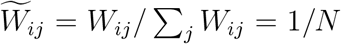. Thus the activation of a node due to the neighboring nodes is simply given by the density of active nodes in the network, i.e., the argument inside Θ[·] in (2) becomes

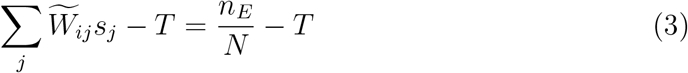

and it is independent of the particular node *i*, i.e., *r*_act_(*i*) = *r*_act_.

These transition rules induce a Markovian dynamics on *n_E_*, *n_R_*, *n_Q_* = *N* − *n_E_* − *n_Q_*, respectively the number of active, refractory, and inactive nodes, with the following rates:

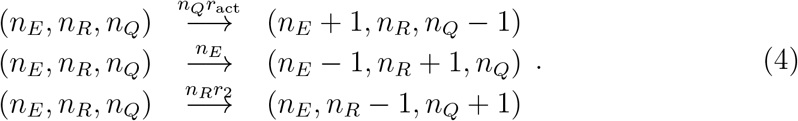

Then, from the reactions in (4), we can write the master equation of our continuous-time model

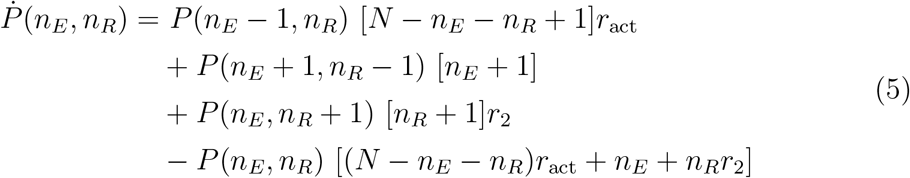

where *P*(*n_E_, n_R_*) is the joint probability of *n_E_* active nodes and *n_R_* refractory nodes.

### 2.2. Equilibria and power spectrum

In order to study analytically the dynamics given by the master equation (5), we perform its Kramers-Moyal expansion truncated at second order. In this way, we can derive the associated Fokker-Planck and Langevin equations [38]. The latter describes the stochastic evolution of the density of active *x* = *n_E_*/*N* and refractory *y* = *n_R_/N* nodes, which obeys

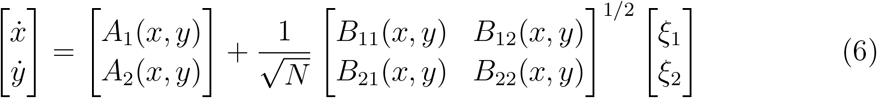

where ***ξ*** = [*ξ*_1_, *ξ*_2_] is an uncorrelated 2d white Gaussian noise, i.e., such that *ξ_i_* ~ *N*(0, 1) and 〈*ξ_i_*(*t*)*ξ_j_*(*t*′)) = *δ_ij_δ*(*t* − *t′*), ***A***(*x, y*) is the deterministic drift term, and ***B***(*x, y*) encloses the stochastic diffusive part (see Appendix A for the full derivation).

To analytically investigate the oscillatory dynamics of our model, from (6) we perform a linear noise approximation [38] by defining the local coordinates (*ζ*_1_, *ζ*_2_) as

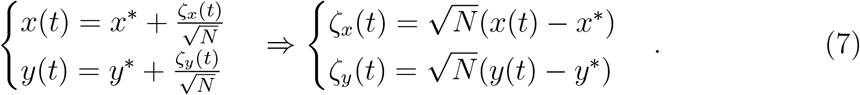

Then (see Appendix C for details) the power spectrum of the oscillations around a given equilibrium is given by

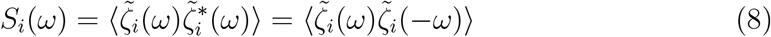

for *i* = *x, y*.

To test the validity of our analytical predictions, we simulate the dynamics of the continuous model with a discretization step of Δ*t* = 0.01 starting from a random configuration of active and refractory neurons. The parameters *r*_1_ = 0.001 and *r*_2_ = 0.1 remain fixed in all simulations and are chosen to be similar to the ones used in previous works [16, 18, 34, 35, 36, 37].

## 3. Results

### 3.1. Existence of a bistable region

In the limit of a large number of interacting units in the system, the effect of random fluctuations becomes negligible. In fact, in the thermodynamic limit, *N* → ∞, the time evolution of the densities described by (6) converge, over a finite time interval, to the solutions of the following system of differential equations

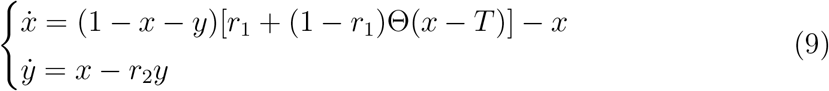

which describes the deterministic evolution of the density of active and refractory units. Although we cannot obtain the full analytical solution of (9), we can study the system’s equilibria and their stability. Indeed, by varying the threshold *T* the dynamics switches between two different regimes based on the value of Θ[·], as we see in figure 1. These two phases are characterized by high and low levels of activity, respectively. We call them super- and sub-critical phase.

**Figure 1:**
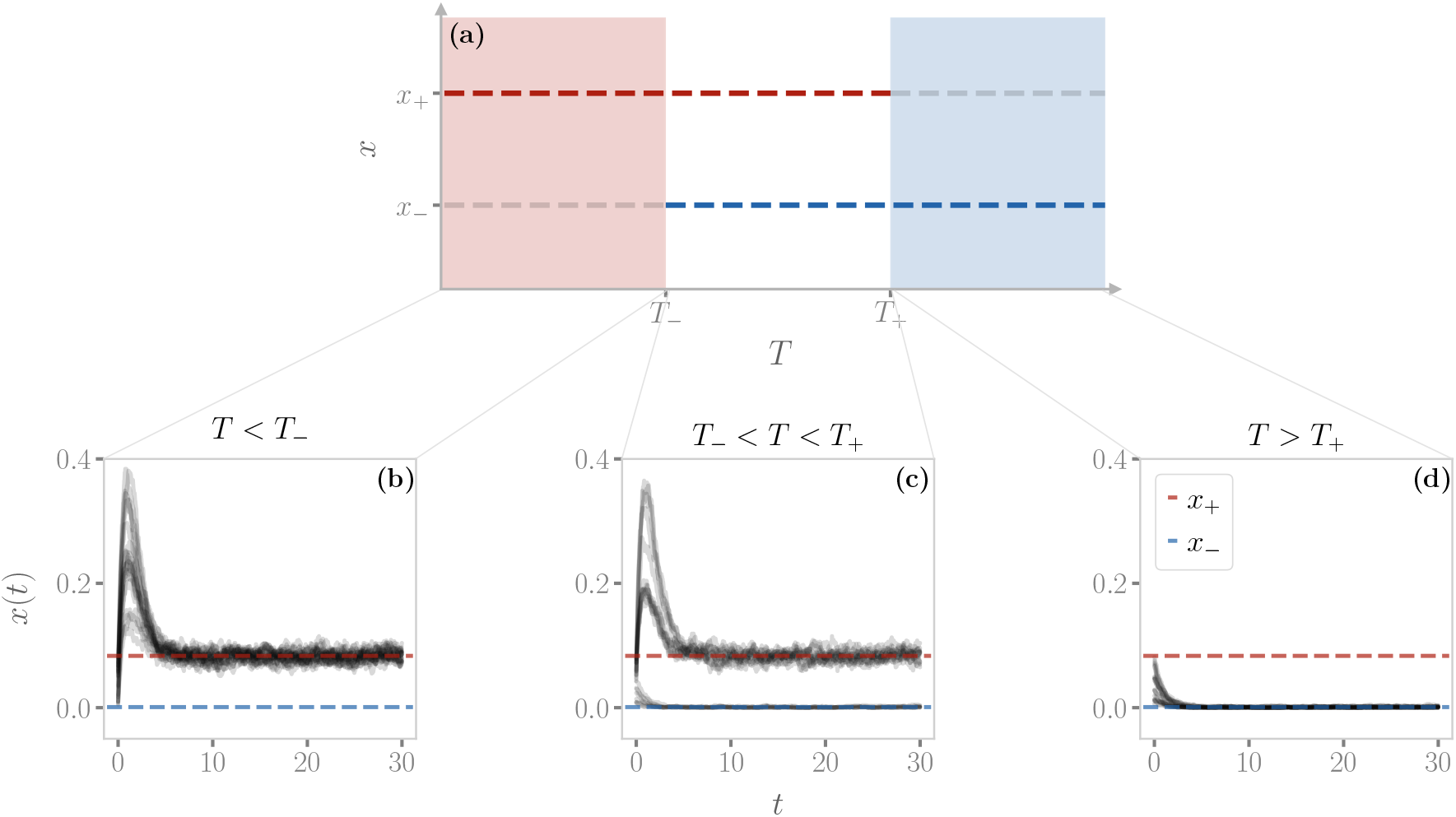
Diagram of equilibria in the model. (a) The region of existence of super- and sub-critical equilibria. As the control parameter *T* changes, we can identify three different regions: for low *T*, only the supercritical equilibrium exists (*red region*); for high *T*, only the subcritical equilibrium exists (*blue region*); for intermediate values of *T*, the two equilibria coexist. (b-d) Examples of trajectories in the three regions. Model simulated with a fully-connected network of size *N* = 10^3^. Each plot shows 30 trajectories from a random initial configuration.

The super-critical phase is defined by the condition *x* > *T*, for which the Heaviside function in (9) evaluates to 1. Hence, and at stationarity we find

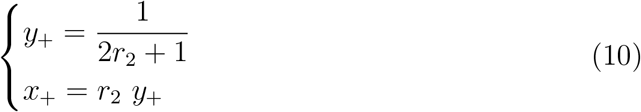

so that in this regime the average activity *x*_+_ is independent of the rate of self-activation *r*_1_. This means that the spreading of the activity is completely driven by the interaction between active neighbors. For this equilibrium to exist, we need

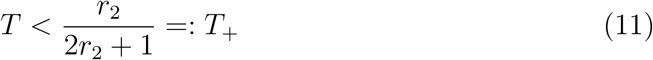

so that the inequality *x* > *T* is satisfied. This defines the threshold below which the super-critical phase exists.

Likewise, the sub-critical phase is defined by *x* ≤ *T*. At stationary, (9) leads to

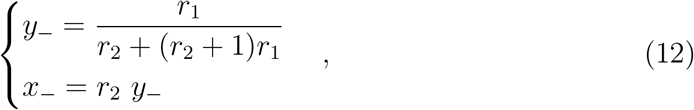

and the inequality *x* ≤ *T* implies that

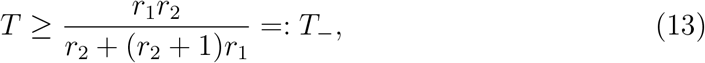

i.e., above the threshold *T*_−_ the sub-critical phase exists. As expected, from (10) and (12) we notice that ∀*r*_1_, *r*_2_ the fraction of active nodes *x*_+_ in the supercritical phase is larger than the subcritical equilibrium *x*_−_, since *r*_1_ ≤ 1. Moreover, in the range of *T* given by equations (11) and (13) for which such solutions exist, they are both stable equilibria, each with its own basin of attraction (see Appendix B for the extensive analysis).

Crucially, and ∀*r*_1_, *r*_2_, (11) and (13) imply that *T*_−_ < *T*_+_, thus three regions emerge in the parameter space spanned by *T*, as shown in Figure 1. For *T* ≤ *T*_−_, the sub-critical equilibrium does not exist, hence we can only observe the super-critical equilibrium. On the other hand, for *T* > *T*_+_ only the sub-critical equilibrium exists. In between these values, for *T*_−_ < *T* ≤ *T*_+_, the two equilibria coexist and we find a region of bistability.

### 3.2. Power spectrum

Neural activity typically exhibits a certain level of stochastic fluctuations, even when the brain is at rest. In fact, a growing amount of evidence suggests that neural noise might enhance the signal processing capabilities of neurons [39, 40]. To this end, we explore analytically the presence of oscillations in the model through the stochastic linearization obtained via a system size expansion [38], from which we derive the temporal evolution of the fluctuations around the equilibria (see Appendix C). Indeed, this approach has proven to be effective in other neuronal models [27, 41, 42, 43]. We find that the power spectrum in the super-critical regime is given by

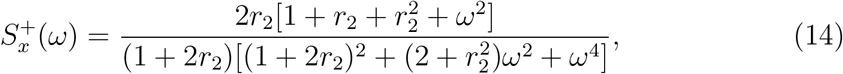

and in the sub-critical regime we have

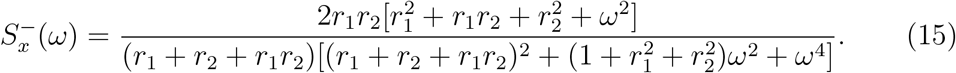

Power spectra obtained from the simulation of the model are perfectly matched by these theoretical expressions, as we see in figure 2.

**Figure 2:**
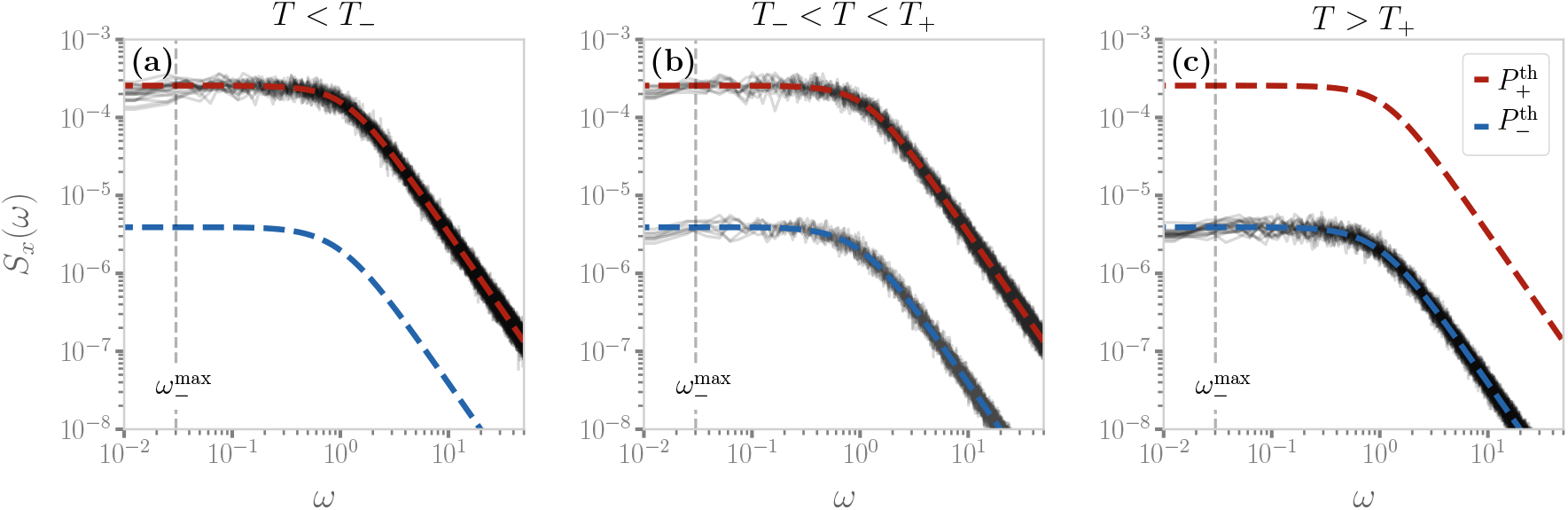
Theoretical expressions of the power spectrum are well-matched by simulated data from a fully-connected network at stationarity. The gray lines represent the power spectrum obtained by simulating the continuous-time model for 10^5^ steps at stationarity in a network of *N* = 10^3^ nodes and after an initial transient of 5 · 10^4^ steps. (a) For *T* < *T*_−_ the power spectrum shows a small peak at 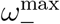. (b) In the bistable region, depending on which equilibria the dynamics settles, we can find both power spectra. (c) For *T* > *T*_+_, no peak emerges in the power spectrum.

Equations (14) and (15) show that, in both regimes, for low frequencies the power spectrum is flat. On the other hand, in the large frequencies limit, we find Brownian noise, i.e., *S*(*ω*) ≈ *ω*^−2^. Such scale-free behavior of the frequencies’ spectrum is found, for instance, in Local Field Potentials (LFPs), i.e., the electrical activity of the brain measured with single microelectrodes [44].

Notably, in the super-critical regime, (15) does not display any peak. A small peak at

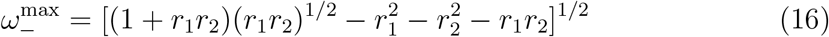

emerges instead in the sub-critical phase (15) for the range of parameters in which 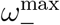 exists, i.e., the radical is non negative. These results suggests that in the mean-field limit of the model, stochastic amplifications alone are not sufficient to induce significant sustained collective oscillations.

### 3.3. Finite-size effects

In order to assess the effects of finite sizes on the region of bistability, we track the average activity 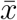 as an order parameter following the approach used in [34]. The simulation starts at *T*_0_ = 0.2 · *T*_−_ from a random initial configuration and, after a given number of steps, we increase the control parameter *T* by a small Δ*T* without resetting the system states. Such procedure is repeated up to a final value *T_F_* = 5·*T*_+_. Then, the same procedure is repeated by starting from *T_F_* and decreasing *T* down to *T*_0_. By doing so, if Δ*T* is small enough, in the coexistence region we should find a hysteresis cycle as a consequence of bistability. Since we want to properly span both the super- and sub-critical regions, and because of the different order of magnitude the two theoretical thresholds (*T*_−_ ≈ 10^−3^, *T*_+_ ≈ 10^−1^), we choose to take 60 logarithmic steps.

In figure 3(a) we plot the behavior of 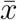 at different steps of this procedure for fully-connected topologies with different sizes. In the super- and sub-critical region, 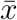 is in accordance with the theoretical predictions (10) and (12). In between the theoretical values of *T*_±_, we recover the discontinuous transition and the hysteresis cycle previously found in [34, 36]. Perhaps unsurprisingly, for small network sizes the limits of the hysteresis cycle do not precisely match the expected values of *T*_+_ and *T*_−_ given in (11) and (13). In fact, due to the finite size of the system, the associated noise contribution causes the bistable region to shrink as the size of the network is reduced.

**Figure 3:**
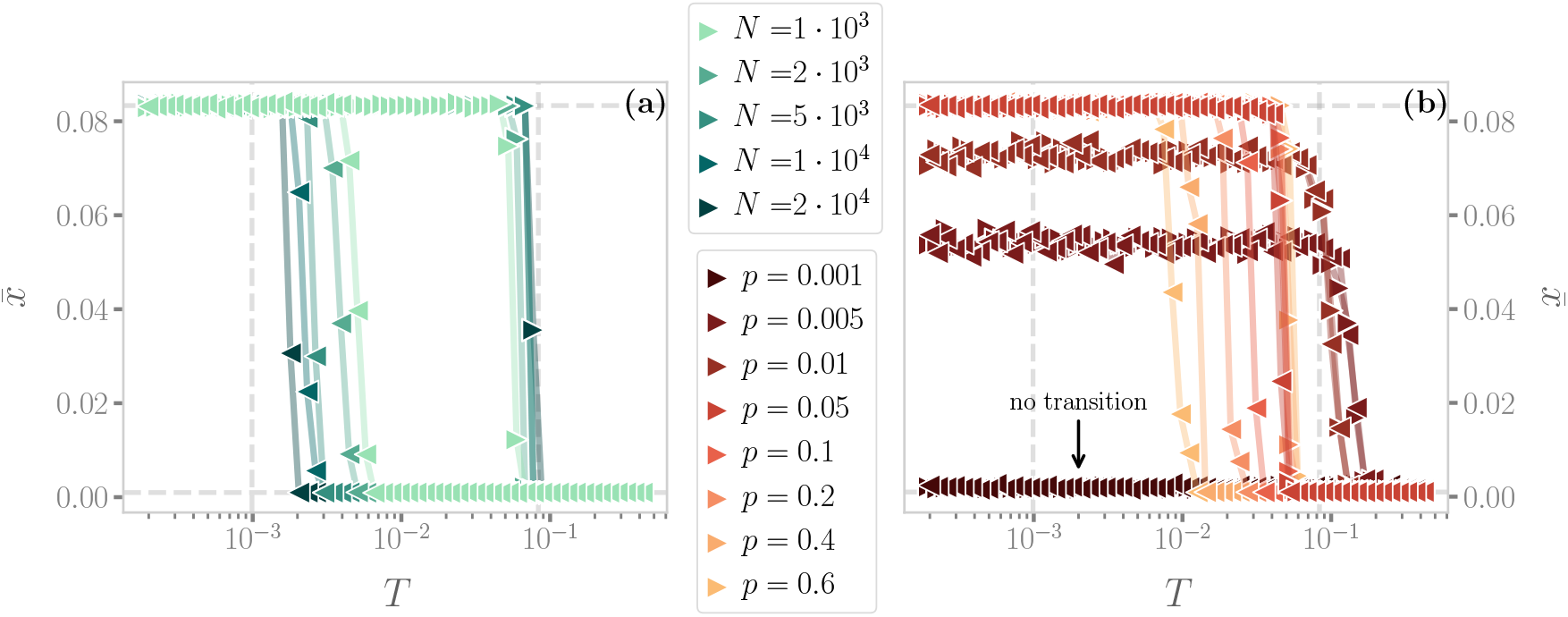
Average activity 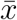 as a function of the control parameter *T* for different sizes and topologies. To probe the presence of the bistable region, we slowly change *T* every 10^5^ steps starting from *T*_0_ = 0.2 · *T*_−_ up to *T_F_* = 5 · *T*_+_ for 60 values of T taken in a logarithmic scale, and then decreasing it back to *T*_0_ in the same way. (a) Results obtained with fully-connected topologies of different sizes N. Triangles to the right represent values obtained, while increasing *T*, whereas triangles to the left represent values obtained while decreasing it. We see that an hysteresis cycle emerges, and that it approaches the expected boundaries of the bistable region as *N* increases. (b) Results obtained from Erdős-Rényi networks with *N* = 10^3^, constant weights and for different wiring probabilities *p*. As the connectivity decreases, the bistable region shrinks and the transition between the two regimes becomes smooth. For extremely low values of *p*, the transition disappears (as indicated by the arrow) and the system is never in the super-critical regime.

### 3.4. Effects of the network structure

Insofar, we have considered the mean-field limit only, which corresponds to a fully-connected topology with constant weights. However, the architecture of the brain is usually characterized by sparse connectivity, and brain networks often display a non-trivial topology with small-world properties and community structures, both at the micro- and macro-scale. Moreover, the strength of the interaction between different brain regions is highly heterogeneous and typically follows a scale-free distribution [32, 45]. Hence, although eased by the homeostatic normalization [16], the hypothesis of constant weights is not fulfilled as well. In this section, we relax the first of these assumptions - a fully connected topology - at the price of analytical tractability.

We first study the simple case of an Erdős-Rényi network with a given wiring probability *p* between two nodes and constant weights [46]. We repeat the procedure described in section 3.3 at fixed network size, but for different wiring probabilities, see figure 3b. We find that, as we lower the connectivity, the bistable region shrinks until it disappears, giving rise to a smooth transition at low values of *p*. This behavior, which is deeply different from the one expected from the mean-field approximation, is consistent with previous results obtained in the discrete time model [34, 37]. In the next section we show that such smooth transitions are strengthened by the introduction of empirical connectivity and they are crucial for the onset of emergent collective oscillations. Eventually, for very low values of *p*, the transition disappears as it becomes impossible for the network to sustain the super-critical regime..

### 3.5. Emergence of collective oscillations and continuous transitions

We now consider an empirical connectome of the human cerebral cortex with *N* = 998 regions [32]. In this case, we have both a complex topology and a non-trivial distribution of weights, as we see in figure 4a. We find that, quite surprisingly, numerical simulations show that the analytical expressions of the two equilibria are still valid in the limit of small and large values of *T*. However, for intermediate values of the control parameter the average activity is no longer bounded to the two equilibria, but it rather changes continuously from one to the other, as we see in figure 4b.

**Figure 4:**
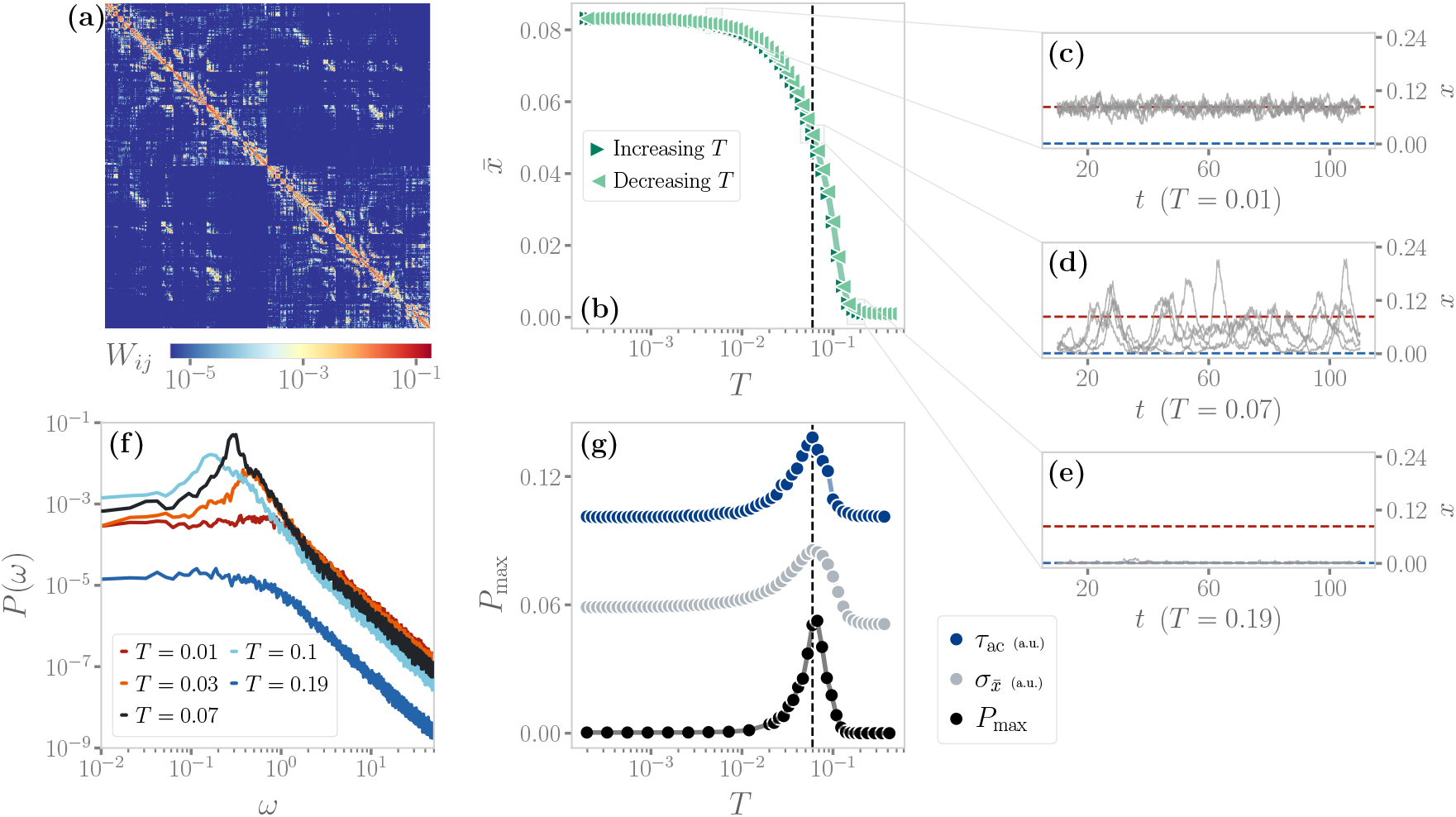
Dynamics of the model over an empirical connectome shows the emergence of a critical-like transition and collective oscillations. (a) Plot of the connectome from [32] (b) The average activity 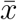 does not show any hysteresis cycle but rather changes smoothly from *x*_+_ to *x*_−_ as *T* increases. (c-e) Examples of trajectories at different values of *T*. In particular, at intermediate values of *T*, the trajectories show high variability and a rich dynamics. (f) The power spectrum at different values of *T*. As 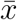 smoothly changes between the two equilibria, collective oscillations emerge. Notice that the frequency peak is at a frequency higher than 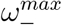. (g) The peak of the power-spectrum *P*_max_ is maximal at intermediate values of *T*. At the same point, both the autocorrelation time *τ*_ac_ and the variance 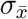 of 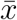 peak (shown in arbitrary units), suggesting that a critical-like transition might be present.

In figures 4c-d-e we plot some trajectories for different values of *T*. We clearly see that, at intermediate values of *T*, the bistability is not present anymore - signaling that the transition is not first-order anymore, but rather happens in a continuous fashion. In particular, in figure 4f we show that the power spectrum now has a peak at these intermediate values - i.e., collective oscillations emerge. Crucially, the value of such peak *P*_max_ is maximal at intermediate values of *T*, where the average activity 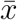 is in between the equilibria *x*_±_. In figure 4g we show that at this particular value of *T* the variance of the activity 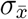 peaks as well. Moreover, the activity shows long-range temporal correlations - indeed, the autocorrelation time *τ*_ac_, computed as the characteristic decay time of the autocorrelation function, peaks at the same value of *T*.

These features are typically found in finite-size systems close to a second-order phase transition [47, 48], suggesting that they may emerge from a critical point of the control parameter *T*. Hence, the transition observed in the presence of the empirical connectome is reminiscent of criticality, rather than the bistability predicted by the mean-field limit. Let us stress that these features are emerging at the dynamical level, contrary to the percolation transition originally studied by Haimovici *et al*. [18]. In general, we find that this dynamical transition does not happen at the same value of *T* of the percolation transition, as observed in other models [33].

In order to understand the relevance of the non-trivial topology of the connectome, we consider the Erdős-Rényi network previously studies, but with the same wiring probability of the connectome, *p*_conn_ ≈ 0. 08, and both with and without weights re-sampled from the empirical connectome weight distribution. As expected from figure 3, without the weights the wiring probability is high enough that the transition is discontinuous and a bistable region exists. In figure 5a-e we see that, in this scenario, the null model matches the behavior of the mean-field limit. No peak in the power spectrum emerges, and the stationary dynamics always reaches one of the two equilibria *x*_±_.

**Figure 5:**
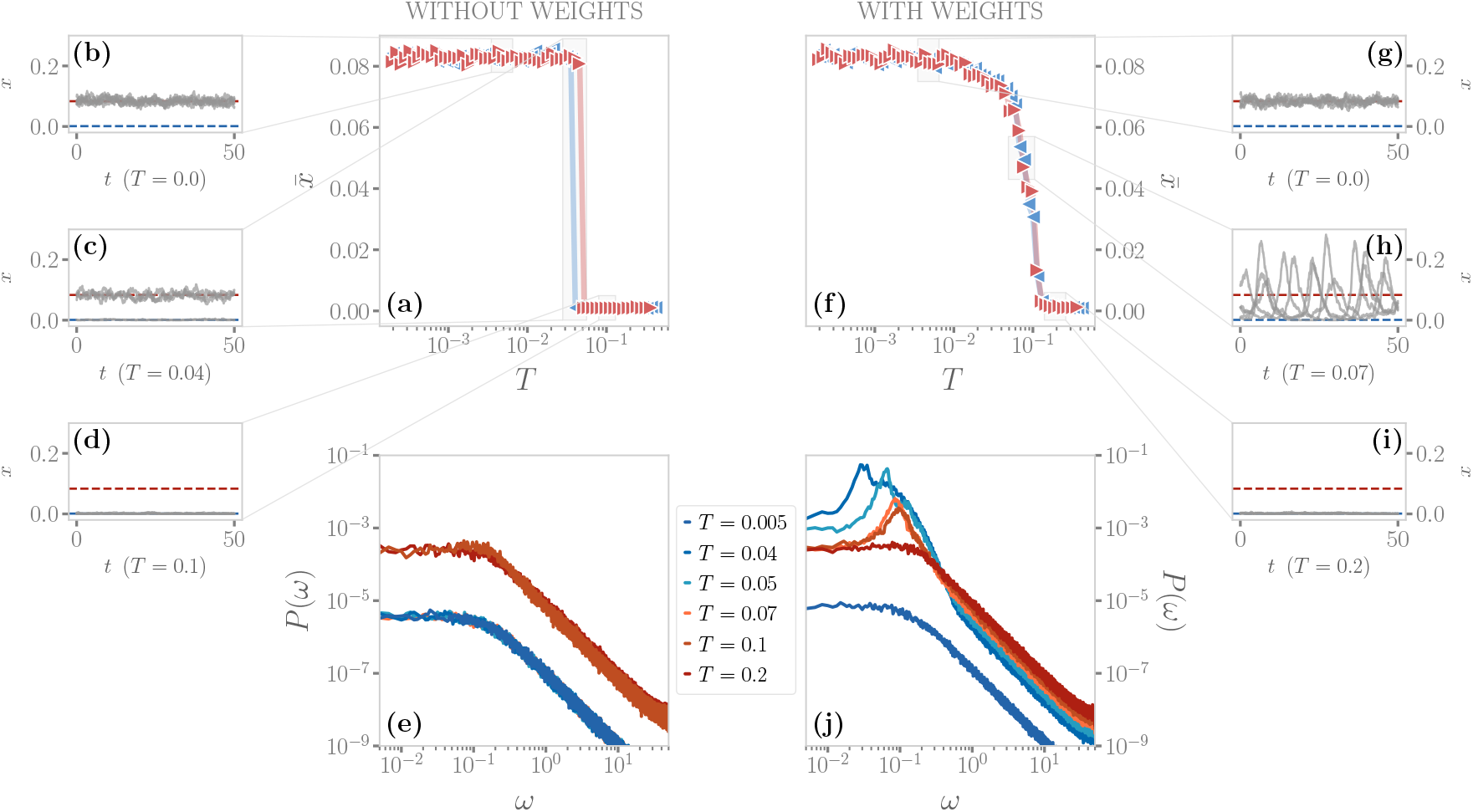
Dynamics of the model over an Erdős-Rényi network with wiring probability *p*_conn_ ≈ 0.08, without and with weights following the empirical distribution of the connectome. (a-d) Without the weights, the transition is discontinuous with a bistable region, as we can see from both the hysteresis cycle and the trajectories. (e) No oscillations are present, and the power spectrum follows the mean-field prediction. (f-i) In the presence of the empirical weights, the transition becomes instead continuous. The trajectories in the region where the average activity 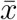 changes smoothly between *x*_±_ display large departures from the expected equilibria. (j) With the empirical weights, oscillations are present.

These results change dramatically when we add back the weights from the empirical connectome. In figure 5f-i we see that no hysteresis cycle emerges, as the transition is now continuous, and the system displays an oscillating behavior. Indeed, the power spectrum in figure 5j displays a clear peak, as we have previously shown for the connectome. That is, the presence of the empirical weights helps the disruption of the bistable region predicted at the mean-field level. Importantly, in Appendix D we show that such disruption emerges only if the wiring probability *p* of the Erdős-Rényi network is low enough. In fact, even with the empirical weights, the bistable region is still present in a fully connected network. This strongly suggests that a continuous, critical-like dynamical transition with global oscillations emerges if the underlying network is either extremely sparse - as in figure 3 - or at higher values of *p*, but with a heterogeneous weights distribution. Crucially, empirical connectomes are often characterized by such features.

## 4. Discussion

Models of large-scale neuronal dynamics are fundamental in explaining and predicting neuronal activity at the macroscopic scale [30]. Such models, describing the collective behavior of populations of neurons and biophysically-inspired, often replicate observed patterns of brain dynamics, e.g., scale-free avalanches [10, 15, 49], long-range correlations [18, 19], global oscillations [25, 26, 27]. However, the collective dynamics is crucially determined both by the dynamical rules that model the inter-neuronal activations and by the geometry of their connections [34]. Furthermore, shared modulations of neural activity may play an important role and unexpectedly explain some of these properties [19, 50]. Disentangling such distinct contributions is a fundamental step to gaining a deeper and more explicit understanding of the mechanisms behind the emergent patterns observed in the brain.

Driven by these considerations, in our work we have developed a continuous-time version of a whole-brain stochastic model [18] and we have studied the nature of the associated critical transition. Previous efforts were often focused on the percolation transition emerging in the model, discussing the effect of the topology in shaping the transition by means of in-silico [35, 36] and empirical connectomes [16, 37]. Here, we focused instead on the dynamical transition arising in this model [34]. To our current knowledge, this study is the first attempt to investigate the nature and the consequences of this dynamical transition from an analytical perspective.

Yet, the bistable region found in the mean-field limit lacks any sign of collective oscillations. However, we have shown that this bistability can be disrupted by an interplay between the underlying network sparsity and a heterogeneous enough weight distribution. Crucially, these properties are typically found in empirical connectomes of the human brain. In this scenario, the bistable region vanishes and a continuous critical-like transition emerges, with large autocorrelation times and variability of neural activity. At this transition, we also observe large collective oscillations, suggesting that both criticality and network structure play a fundamental role in driving the collective behavior of neurons.

Importantly, we can also compare the trajectories in the empirical connectome and in the null random netwrk model with weights, see figure 6. In the empirical connectome, the dynamics is typically richer and oscillates close to the nullcline 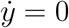 predicted by the mean-field (9). On the other hand, in the Erdős–Rényi case, the trajectories display large transient dynamics away from the (*y*_−_, *x*_−_) equilibrium, reminiscent of noise-induced oscillations [27, 41, 42, 43] or non-normal systems [51, 52]. This suggests that, although both models display emerging oscillations, the underlying dynamical features might be different. Notably, these phenomena have usually been observed in models with excitatory and inhibitory populations. Here, we rather have a single excitatory population with a refractory state, hinting that the two different scenario may lead to a similar phenomenology. Therefore, further work is needed to explore the role of higher-order structures in the empirical connectome - e.g., modularity [53, 54] or heterogeneity [55] in the degree distribution - and their effect on the model dynamics.

**Figure 6:**
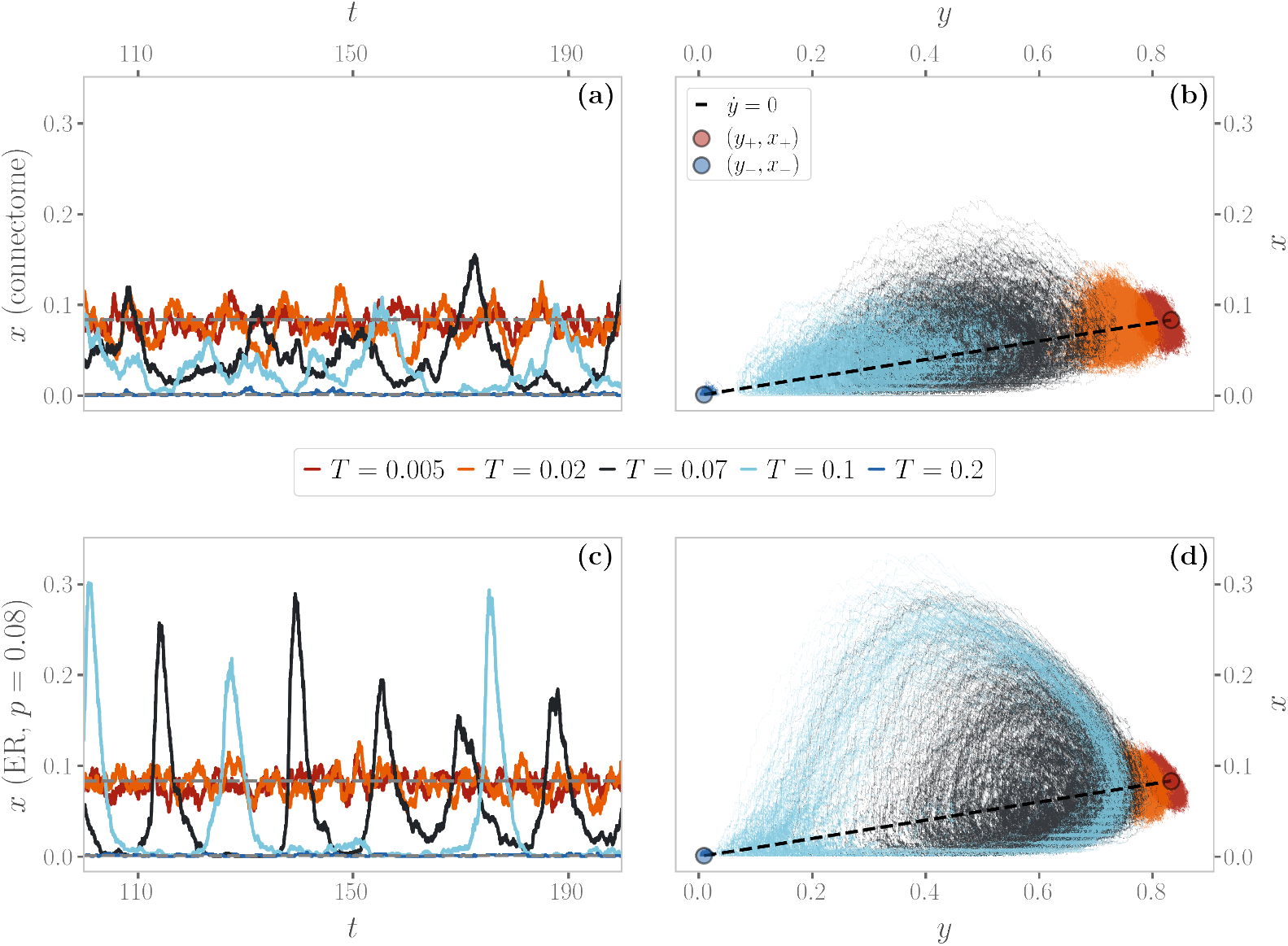
Comparison of the trajectories in the empirical connectome and in the Erdős-Rényi with *p* = *p*_conn_ and empirical weights. (a) Connectome trajectories of *x* as a function of time and (b) in the phase space for different *T*. The black dashed line in panel (b) is the nullcline of the refractory population *y*, the gray lines in panel (a) are *x*_±_. (c-d) Same, but in the case of the null model. Notice how the trajectories in the phase space are qualitatively different, and oscillations emerge as large activations from the sub-critical equilibrium (*y*_−_, *x*_−_).

The small size of the empirical connectome considered here may be a limitation to these investigations, since finite-size corrections may be hiding criticality. Notably, in [56] a similar modification of the discrete-time Greenberg & Hastings model run on a large-scale connectome displays semicritical behaviors consistent with a Griffith phase in a certain range of the control parameter. Such use of synthetic connectomes overcomes the finite size issue, at the cost of relying on some subjective assumptions about the generated topologies. Hence, future works should be devoted to fully understand whether the observed continuous transition is associated with a real critical point or with other phenomena such as rare region effects [57] or noise-induced transitions [43].

Overall, here we have shown in detail how network structure plays a fundamental, yet sometimes poorly understood, role. Therefore, we believe that our work will serve as a baseline for future analytical efforts in explaining the nature of the observed transition under more relaxed assumptions, e.g., in presence of a non-trivial distribution of weights and different topologies, to further understand the influence of both in the emergence of critical features in the human brain. Possible approaches may include the use of heterogeneous mean-field methods as done in the study of epidemic spreading [55] or annealed network approximations [58]. All in all, we believe that our findings are a further contribution to the still puzzling “critical brain hypothesis”.

## Acknowledgments

S.S. acknowledges DFA and UNIPD for SUWE BIRD2020 01 grant, and INFN for LINCOLN grant. We acknowledge Dr. Rodrigo Rocha for pointing us to relevant literature. S.S. designed the study, S.S and M.F. supervised the research. G.B. and G.N. performed the analytical calculations, implemented the simulations and prepared the figures. G.B., G.N., B.M., M.F. and S.S. interpreted the results and wrote the manuscript. All authors contributed to the article and approved the submitted version.

## Code availability

The code of HTC model is available at https://github.com/gbarzon/HTC_Mesoscopic_Brain.

## Appendix A. System size expansion in the mean-field approximation

The master equation (5) can be reframed in terms of the density of active *x* and refractory *y* neurons. Since Δ*x* = 1/*N*, Δ*y* = 1/*N*, we can treat them as continuous variables in the limit of a large system, i.e., *N* → ∞, thus *P*(*x, y*) becomes differentiable. By taking the continuum limit of the master equation (5) and by expanding all the terms, i.e., the Kramers-Moyal expansion [38], up to the second order, we obtain the so-called Fokker-Plank equation for the probability density *p*(*x, y*)

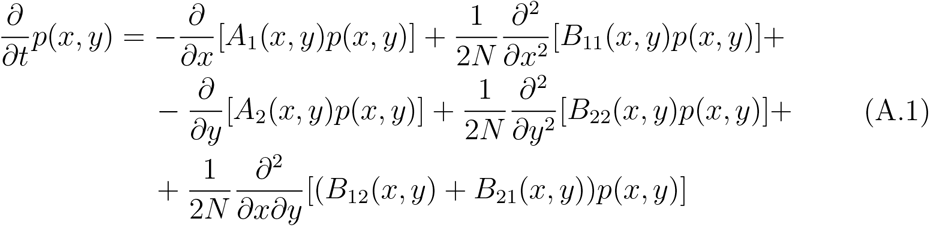

where the coefficients are

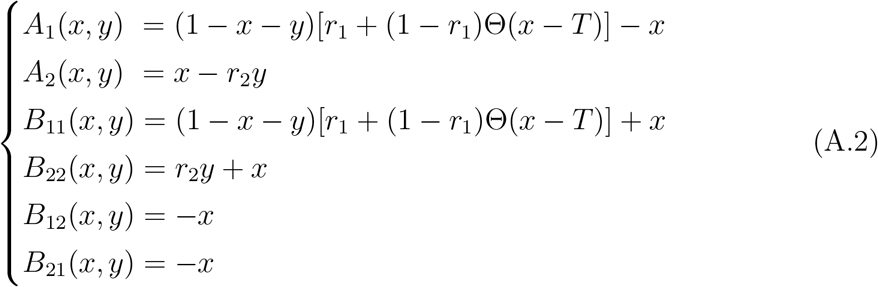

and *P*(*x, y*) = *p*(*x, y*) Δ*x*Δ*y*. The Fokker-Planck equation is a deterministic differential equation describing how the probability distribution of states *p*(*x, y*) evolves over time. Physically, it describes the evolution of an ensemble of systems: if we simulate a huge number of populations of neurons, all with the same parameters, they will have a different evolution due to random fluctuations, but the fraction of systems that have a density of states in [*x, x* + *dx*; *y, y* + *dy*] at time *t* will be given exactly by *p*(*x, y*)*dxdy* (in the limit of an infinite ensemble). An equivalent description can be derived by instead following a single population of neurons. In this case, a change in population density [*dx, dy*] under the effect of stochastic fluctuations *ξ* is given by the associated Langevin equation (6) [38].

## Appendix B. Stability analysis of equilibria

We further investigate the nature of the equilibria through linear stability analysis techniques. Indeed (9) is a dynamical system of the type:

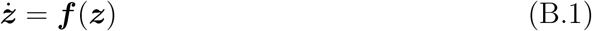

with ***z*** = (*x, y*) a 2-dimensional vector. The equilibria ***z***\* of this system are the ones that satisfy ***f*** (***z***\*) = 0. If we focus on the dynamics near the fixed points, we can perform a change of variables *x* = *x*\* + Δ*x, y* = *y*\* + Δ*y*. In the limit of small variations |Δ***z***| → 0, meaning that we are considering states infinitesimally near the fixed points, (B.1) can be Taylor-expanded as

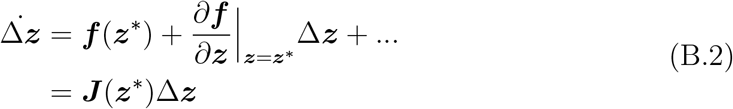

Thus the dynamics near the fixed points is governed, at the first order, only by the Jacobian matrix ***J***. In particular, the (real part of) the eigenvalues *λ* of ***J*** can tell us information regarding the stability or instability. If *max Re*(*λ*) > 0, the trajectories asymptotically diverge from the equilibria, otherwise for *max Re*(*λ*) < 0 the trajectories converge to the fixed point, which is stable in this case.

In the super-critical phase the jacobian evaluated at (*x*_+_, *y*_+_) is

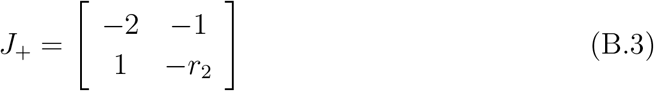

The eigenvalues of (B.3) are

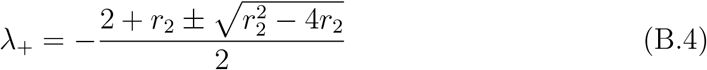

The stability condition holds if *Re*(*λ*_+_) < 0. We can distinguish two regimes: if *r*_2_ ≥ 4 the eigenvalues are purely real, otherwise they have an imaginary part. In both cases the conditions is satisfied, thus the super-critical fixed point (*x*_+_, *y*_+_) is respectively a *stable knot* and *stable focus* (figure B1).

Instead, in the sub-critical phase the jacobian evaluated at [*x*_−_, *y*_−_] is

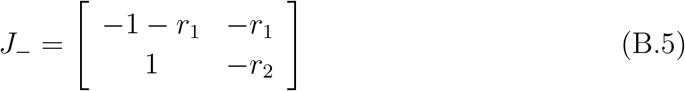

whose eigenvalues are

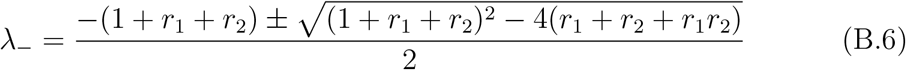

**Figure B1:**
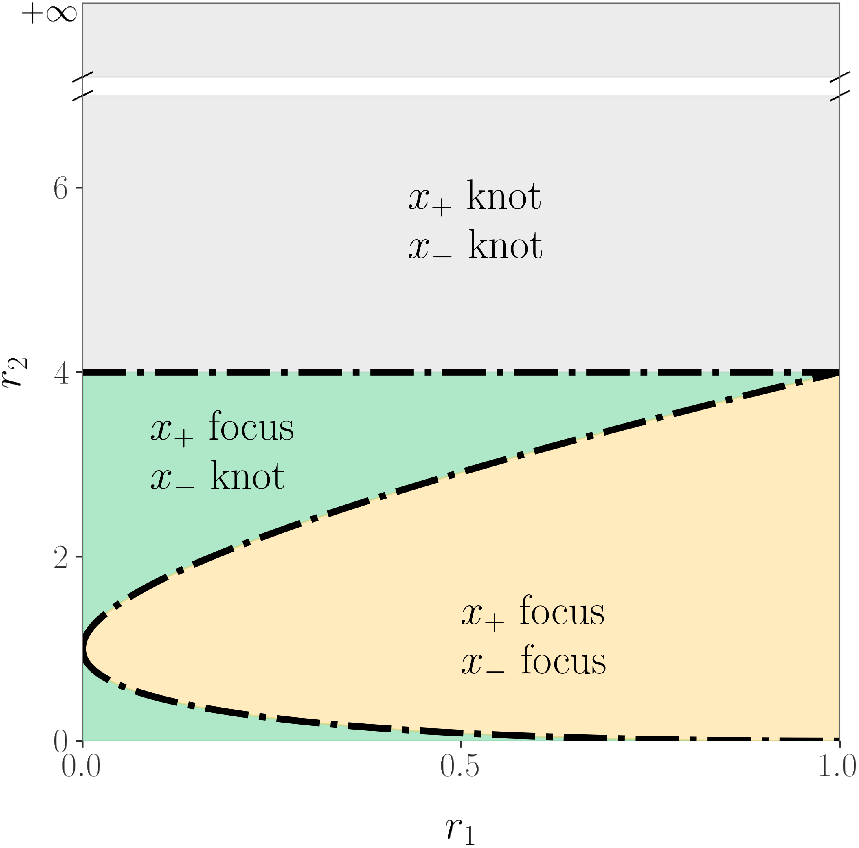
Type of fixed point in the parameter space (*r*_1_, *r*_2_).

The stability condition in this phase is again *Re*(*λ*_−_) < 0, which is satisfied ∀*r*_1_, *r*_2_ (since *r*_1_ ≥ 0 and *r*_2_ ≥ 0). We observe two different regimes by varying the parameters *r*_1_ and *r*_2_: if 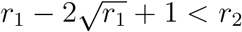 and 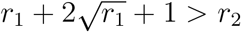 the eigenvalues have an imaginary part, while in the other case they are pure real. So in the first cases the fixed point is a *stable focus*, while it is a *stable knot* in the other case (figure B1).

## Appendix C. Power spectrum

To study the effect of fluctuations around the equilibrium we make use of the linear noise approximation [38]. First, we define here two local coordinates (*ζ*_1_, *ζ*_2_)

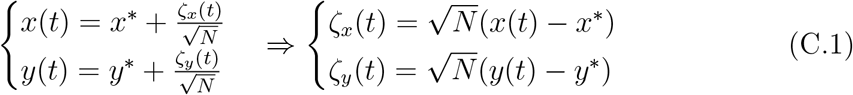

In the Langevin equation, the stochastic fluctuations go as 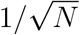, and so here we multiply by 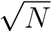 to remove this size dependence.

Then, we rewrite the original equations in terms of (*ζ_x_*, *ζ_y_*), keeping only the linear terms. For the deterministic part, this leaves only the jacobian evaluated at the equilibrium *J*(*x*\*, *y*\*) ≡ *J*. For the diffusion term we need to expand up to 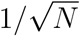 (so then we get order 1 after multiplying by 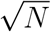). But this means that *b* must be expanded to 0-th order, otherwise we would have terms in 1/*N*, which become 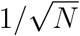 after multiplication, that are negligible in the thermodynamic limit. After this we arrive to:

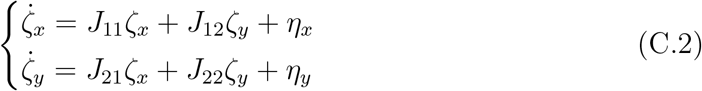

where

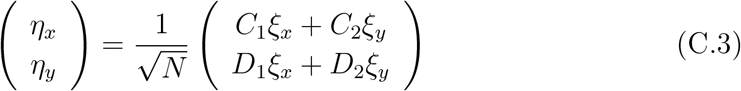

Since *ξ* is Gaussian, also *η* is Gaussian. However, since in *η* we are summing over different components of *ξ*, *η*_1_ and *η*_2_ are not anymore uncorrelated:

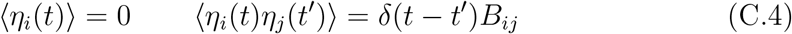

where *B_ij_* = *B_ij_*(*x*\*, *y*\*) is the diffusion matrix evaluated at equilibrium. To proceed, we move to Fourier space. Since the transformation is linear, it preserves the linearity of the equations:

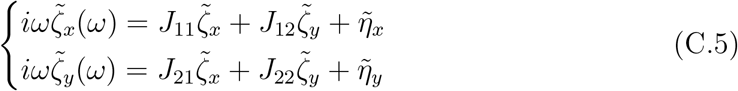

And the statistics of *η* remain the same:

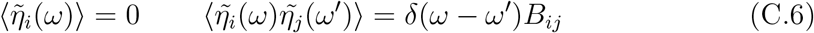

The linear system ((C.5)) leads to:

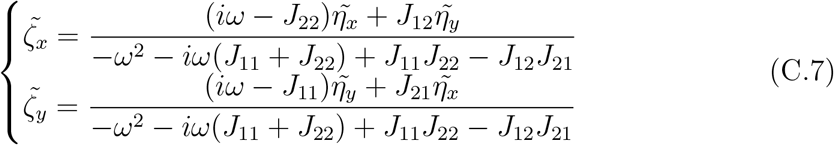

Then we can study the power spectrum

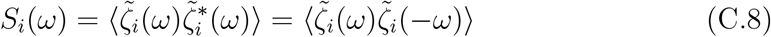

that for the oscillations *ζ_x_* of the density of active neurons *x* leads to

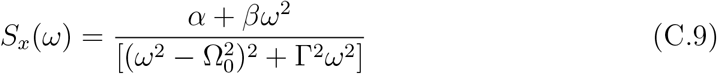

where

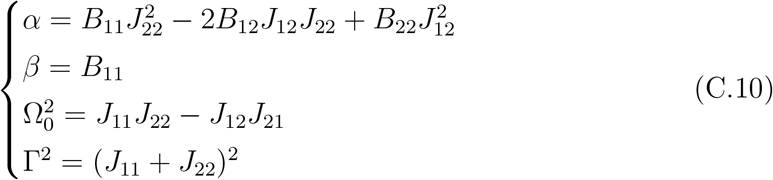

## Appendix D. Effects of empirical weights

Here, we show the effects of the addition of empirical weights to Erdős-Rényi networks. In figure 3 we have shown that with constant unitary weights and the homeostatic approximation, the bistability is still present at high wiring probabilities *p*. The transition only vanishes for very low *p*, and in this scenario the average activity in the super-critical regime is lower than the corresponding mean-field equilibrium *x*_+_. In fact, if the wiring probability is too small, the network is too sparse to sustain activity. Notably, in this regime we find oscillations similar to the ones shown in figure 5h, suggesting that the mechanism at play may be similar. However, the transition is still discontinuous at the wiring probability of the empirical connectome *p*_conn_ ≈ 0.08.

We now add weights re-sampled from the empirical connectome [32], and in figure D1 we show the average activity obtained while slowly varying *T*, as in the main text. First, let us note that in the fully-connected case *p* = 1, even with weights, an hysteresis cycle is still present. Hence, the transition is still discontinuous. However, the transition now becomes smooth already at larger values of *p*, showing that both sparsity and weights aid the disruption of the bistability predicted by the mean-field approximation.

**Figure D1:**
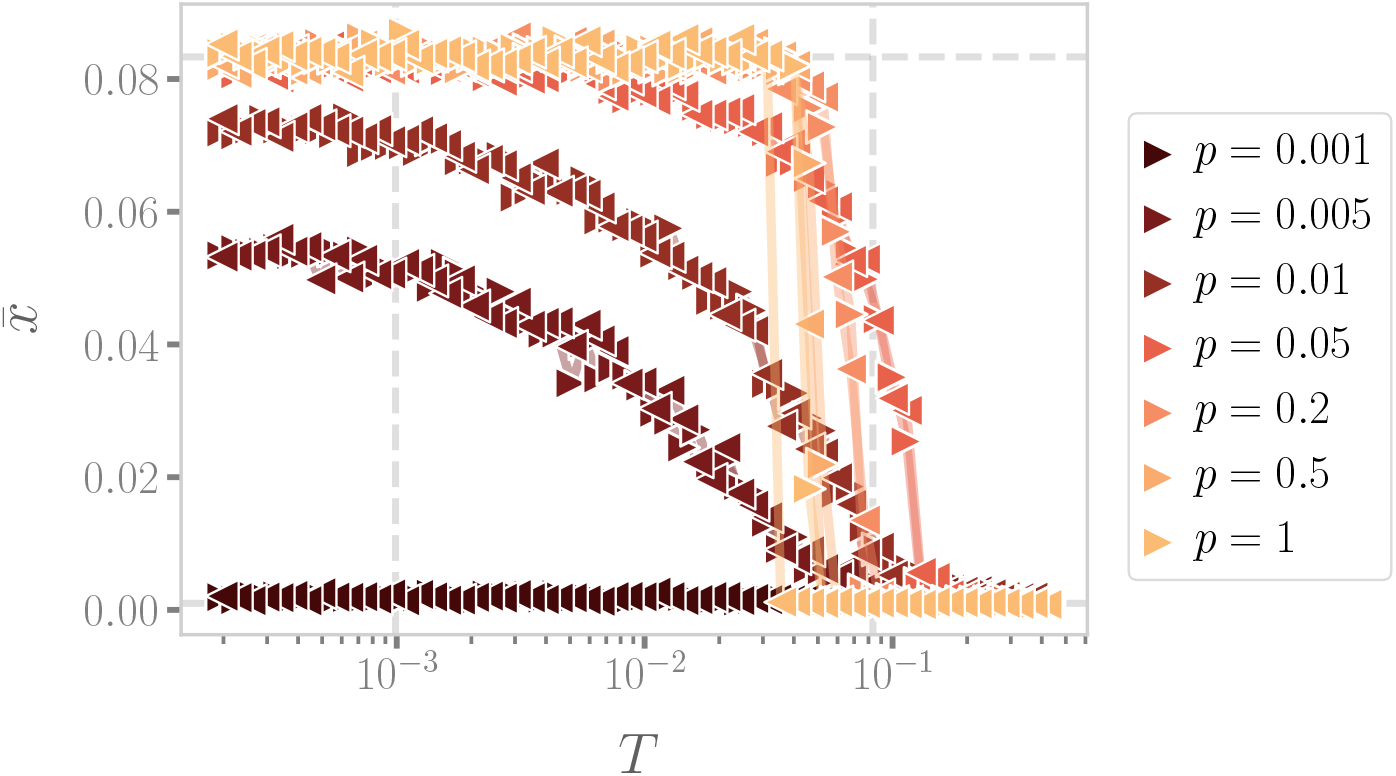
Average activity 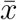 in Erdős-Rényi networks at different wiring probabilities *p*, with weights re-sampled from the empirical connectome. With weights, the transition becomes continuous already at higher *p*.

## Footnotes

‡ We highlight that the parameters *r*_1_ and *r*_2_ in the time discrete model were probabilities, whereas here they are rates.

